# CLAIR: An integrated lipid database across multiple crop species

**DOI:** 10.1101/2023.11.08.566134

**Authors:** Bing He, Mengjia Bu, Qiang Lin, Zhengwei Fu, Junhua Xie, Wei Fan, Jianyang Li, Ruonan Li, Wei Hua, Wanfei Liu, Peng Cui

## Abstract

Oilseed crops, which are rich in plant lipids, provide essential fatty acids for human consumption and serve as major sources of biofuels and essential raw materials for the chemical industry. As a result of population growth and ecological changes, the demand for vegetable oil is increasingly outpacing supply. A comprehensive understanding of the genes involved in lipid metabolism in oilseed crops and the regulatory relationships among these genes is essential for improving oil content. However, current studies on lipid metabolism genes rely heavily on a decade-old database of genes involved in lipid metabolism in *Arabidopsis thaliana*. To address this issue, we mined the literature, integrated data from various databases and studies, and aligned homologs of lipid metabolism genes from nine oilseed crops to construct a comprehensive set of lipid metabolism genes in plants. Using this approach, we identified 221 additional lipid metabolism genes. In addition, we created a user-friendly lipid database called CLAIR (Crop Lipid-Associated Information Resource) by integrating and mining multi-omics data from nine major oil crops. The database is available at http://www.clipair.cn/. These resources should facilitate further research and exploration of lipid metabolism in oil crops, ultimately contributing to improved oil production.

## Introduction

Lipids are an important global commodity and industrial raw material. Most lipids consumed worldwide are derived from plants. In 2017, the total production of vegetable oils and animal fats was estimated to be 206.1 million tons per year, with vegetable oils accounting for ∼87% of this total (Zeb, 2021). The demand for high-quality vegetable oils is expected to double by 2040 as the world’s population continues to grow and industrialize (FAO, 2003). Oilseed crops, a major source of vegetable oils, primarily store triglycerides in seeds and fruits, with variation in the specific types of oils produced (Zanetti et al., 2013). It is imperative to comprehensively understand the genes and mechanisms associated with lipid biosynthesis in different oilseed crops to improve oil content traits in these crops via molecular biology and breeding (Xu and Shanklin, 2016).

Lipid biosynthesis is a complex biological process involving several steps, including *de novo* fatty acid biosynthesis, triglyceride biosynthesis, lipid droplet formation, and lipid degradation (Zafar et al., 2019). This process is closely related to the flux of carbon sources, such as sugar metabolism, and is a typical quantitative trait determined by different gene loci (Rawsthorne, 2002). The major databases for plant lipid metabolism include KEGG (https://www.genome.jp/kegg/pathway.html) (Kanehisa, 2002), MetaCyc (https://metacyc.org/) (Caspi et al., 2020), PlantFA (https://plantfadb.org/) (Ohlrogge et al., 2018), and ARALIP (http://aralip.plantbiology.msu.edu/) (Beisson et al., 2003). The KEGG pathway database relies on manually collating information from scientific literature, where genes, proteins, metabolites, and other components are annotated to create pathway maps. However, since it is not specifically focused on plant lipid metabolism and is primarily designed for visualization purposes, there are certain drawbacks to this database, such as missing annotation information and annotation homogenization (Wrzodek et al., 2013). MetaCyc lists nearly 20,000 metabolites and reactions related to primary and secondary metabolism, making it useful for predicting the metabolic pathways of several metabolites. However, this database emphasizes the presentation of metabolite-related information and does not integrate information on associated genes and proteins (Caspi *et al*., 2020). PlantFA compiles fatty acid composition data from various plants based on literature mining and focuses on summarizing the fatty acid structures of each species in the phylogenetic tree. However, this database does not include pathway annotations or gene information (Ohlrogge *et al*., 2018). ARALIP summarizes and manually annotates more than 600 lipid metabolism-related gene sets and the corresponding pathways in Arabidopsis (*Arabidopsis thaliana*). This database is highly credible and practical and serves as an important reference for functional annotation of oilseed genomes. However, it has not been updated for nearly 10 years (Li-Beisson et al., 2013).

Over the past 10 years, many genes involved in plant lipid metabolism have been discovered (Wan et al., 2020). Despite extensive studies on the regulation of lipid biosynthesis in oilseed crops, most annotations and functional studies of regulatory genes are still based on Arabidopsis homologs or the KEGG database. However, relying on outdated versions of reference gene sets to annotate oilseed genes could lead to errors (Schnoes et al., 2009). To address this issue, here, we reconstructed a reference gene set for lipid metabolism in Arabidopsis. Based on this newly constructed reference, we developed a user-friendly online database called CLAIR (Crop Lipid-Associated Information Resource). This database is dedicated to the lipid metabolism of nine major oilseed species. It incorporates and analyzes multi-omics data from these species to provide a valuable resource and reference for researchers in the field. Using CLAIR, scientists studying lipid metabolism in oilseed crops can access up-to-date, relevant information, reducing the risk of inaccuracies from outdated gene sets. This comprehensive, integrated database is designed to support and enhance research aimed at understanding the regulation of lipid biosynthesis in major oilseed crops.

## Materials and Methods

### Expansion of the lipid metabolism gene set

The Arabidopsis (*Arabidopsis thaliana*) Acyl-Lipid Metabolism gene set, which was published in 2013 as the first lipid metabolism-related gene set in plants (Li-Beisson *et al*., 2013), was used as the starting point to construct our database. To improve and expand this gene set in Arabidopsis, a thorough search of recently published literature on genes associated with lipid metabolism was conducted. Two methods were used to collect relevant literature: searching papers citing the 2013 Arabidopsis dataset and a PubMed search using keywords such as “acyl-lipid”. This comprehensive search returned more than 1200 papers with contents related to lipid metabolism. To evaluate the association of each gene with lipid metabolism, seven levels of evaluation criteria were established. The major criteria included validation by mutation experiments, heterologous expression, and overexpression studies. Validation by KEGG annotation records, literature citations, high similarity to known lipid metabolism-related genes, and any other pertinent evidence was also considered. By applying these seven levels of evaluation, each gene in the gene set was tagged to facilitate further evaluation and screening by researchers. This rigorous, multi-faceted approach was designed to provide an improved and reliable resource for researchers.

### Construction of the CLAIR database

Based on the newly constructed Arabidopsis lipid metabolism gene set, the lipid metabolism genes of nine oilseed crops (rapeseed, soybean, walnut, peanut, sunflower, castor, oil-camellia, oil palm and sesame) were re-annotated by homolog alignment coupled with confirmation using InterPro, UniProt, Gene Ontology Resource, KEGG, and other public databases (Emms and Kelly, 2019; Harris et al., 2004; Kanehisa, 2002; Paysan-Lafosse et al., 2023; UniProt, 2015). To account for the diverse metabolic characteristics of different oilseed crops and to further improve the gene set, genes potentially related to lipid metabolism in these oilseed crops were functionally re-annotated based on their representative genomes, where information from the KEGG database and relevant genes mentioned in the literature were integrated. By carefully examining the genomic data of each oilseed crop, genes likely to be involved in lipid metabolism were identified. To provide a more comprehensive resource, information on major quantitative trait loci (QTLs) and expression QTLs (eQTLs) related to lipid anabolism in each species was also collected. These features make our gene set better tailored to the specific characteristics and metabolic pathways of each oilseed crop (Li et al., 2018; Tam et al., 2019).

After obtaining gene sets related to lipid anabolism in oilseed crops, the CLAIR database was constructed. CLAIR contains information on lipid metabolism genes including gene visualization, gene IDs, associated sequences, gene structures, gene functions, associated pathways, and related references. In addition, to investigate the expression patterns of lipid metabolism-related genes during different stages of seed development, nearly 2000 batches of RNA-seq data from nine oilseed species available in public databases were integrated into our database. To improve the classification of the transcriptome data, all data based on specific processing conditions were manually labeled and a transcripts per million (TPM) expression matrix of all genes was obtained using the alignment-free strategy (Bray et al., 2016). The expression matrix was transformed into specificity measure (SPM) format, with values ranging from 0 to 1, to accurately represent gene expression within each sample. This transformation effectively mitigated the batch effect resulting from different experiments (Julca et al., 2021). In addition, representative transcriptomic data were selected from each species to construct gene co-expression networks and to identify strongly associated modules related to lipid metabolism (Langfelder and Horvath, 2008).

### Database implementation

CLAIR was developed using the programming language PHP, the ThinkPHP 6.1 framework, and a MySQL 8.0 database (http://www.mysql.com; a free, popular relational database management system) for data storage. For the front-end interfaces, the technology stack consists of Vue3 (https://v3.cn.vuejs.org/, an approachable, high-performance, versatile framework for building web user interfaces) and Element UI (https://element.eleme.cn/; a Vue3 component library for designers and developers). Like many other database applications, CLAIR provides user-friendly access through its web interface. CLAIR’s web interface has an intuitive navigation section that provides convenient links to key functionalities and web pages. These include resource collections for gene sets, protein families, QTL information, pathways, expression profiles, tools, downloads, and other informative pages.

#### Contents and usage

##### 1. Integration and display of gene set information

This newly constructed Arabidopsis lipid metabolism gene set contains 996 genes, including 221 newly added genes compared to the original dataset. Importantly, all 221 additional genes are well supported by evidence from the literature. In addition, 112 of these new genes have been annotated by the KEGG database, with the largest subset (31 genes) involved in the sterol biosynthesis-related pathway. Of the 996 genes in the complete set, 679 are supported by literature evidence and 380 have been validated by molecular experiments, further confirming their involvement in lipid metabolism processes (Figure 1B). Notably, our gene set includes 54 lipid-related transcription factor genes that may play critical roles in regulating lipid metabolism. Each lipid metabolism-related gene in our dataset is annotated with essential information about the subcellular localization of its product, the specific pathway in which it functions, the gene’s location, the name of the encoded protein family, the gene name, functional evidence, and related references.

**Figure 1.**
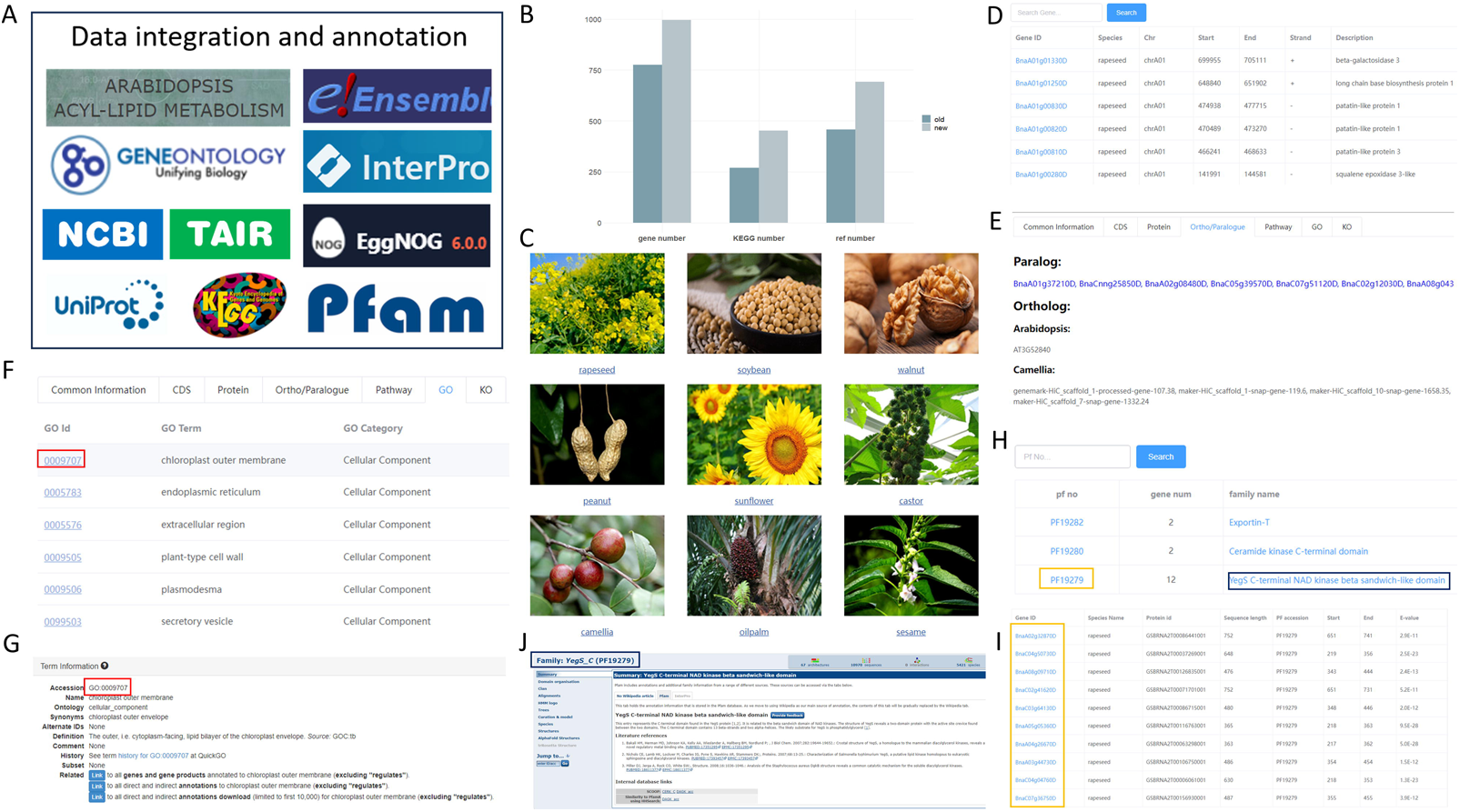
Overview of data processing and gene set contents of CLAIR. (A) General information about the data integration and annotation process. (B) The results of our newly integrated gene set and the original data set were compared in terms of the total number of lipid-related genes (gene number), the number of genes annotated by the KEGG database (KEGG number), and the number of genes supported by literature evidence (ref number). (C) The home page of the CLAIR database. (D) Results of the gene list page displayed after clicking on each species image on the home page. (E) Paralogous and orthologous information for each gene, with each gene ID clickable. (F) GO annotation results for each gene. (G) Term information for each GO ID is displayed when clicked. (H) Summary of protein clustering results in each oilseed species. (I) Results of the gene list included after clicking on a specific protein family ID. (J) Click on a specific protein family to jump to the detailed annotation information.

From the CLAIR main page, users can access the summary page of lipid metabolism-related resources for each oilseed species by clicking on the respective species image (Figure 1C). The summary page presents essential information from the core modules, which include basic details, lipid metabolism gene sets, protein families, pathway information, and co-expression networks of the nine major oilseed crops. Within the gene set module, each lipid metabolism gene displayed in CLAIR is clickable (Figure 1D). The “Common information” section provides details such as the gene name, genome localization, associated transcripts, coding sequences, and protein family. In the “Protein” section, users can find the symbol of the protein encoded by the gene, as well as the SwissProt ID (Boutet et al., 2016) and TrEMBL ID (Bairoch and Apweiler, 2000), along with all associated sequences. The “Ortholog/Paralog” section compiles information on orthologs and paralogs of the gene in the nine oilseed species and Arabidopsis (Figure 1E). Clicking on the homolog ID in Arabidopsis will direct users to the gene’s details page in the TAIR database (https://www.arabidopsis.org/) (Rhee et al., 2003), and clicking on the IDs of other oilseed species will also take the user to the gene information page in CLAIR.

The “Pathway,” “GO”, and “KEGG” sections collectively present functional annotation information related to the gene (Figure 1F). Clicking on both the gene ID and GO number links will lead users to the corresponding pages in related public databases, where they can further explore general information about the gene in the lipid metabolism pathway (Figure 1G). The home page of each oilseed species includes not only the gene set module, but also the protein family and pathway modules, providing users with comprehensive information (Figure 1H). In the protein family module, users can find the IDs, family members, and names of all lipid metabolism-related proteins specific to each species (Figure 1I). In addition, the module provides direct links to the PFAM database (http://pfam-legacy.xfam.org/) for further exploration and detailed information on the protein families (Bateman et al., 2004) (Figure 1J).

The “Pathway” module on the home page displays classification results and the number of related genes involved in metabolic pathways, including fatty acid biosynthesis, fatty acid elongation, fatty acid degradation, and other relevant pathways (Figure 2A). By clicking on the pathway name, the detailed gene information and the EC numbers of enzymes involved in all oil metabolism pathways in oilseed species (Figure 2B). Users can also directly access the corresponding information in the KEGG database by clicking on the pathway numbers (Figure 2C, 2D). The pooled results for genes related to the same pathway in different species allow users to comparatively analyze oil metabolism genes in different oilseed species. Moreover, each gene in the gene set module is supported by genome visualization, enhancing the user’s understanding and analysis of the gene’s characteristics and functions (Figure 2E). By clicking on the relevant region in the browser, further detailed information on the genes in that region can be obtained (Figure 2F).

**Figure 2.**
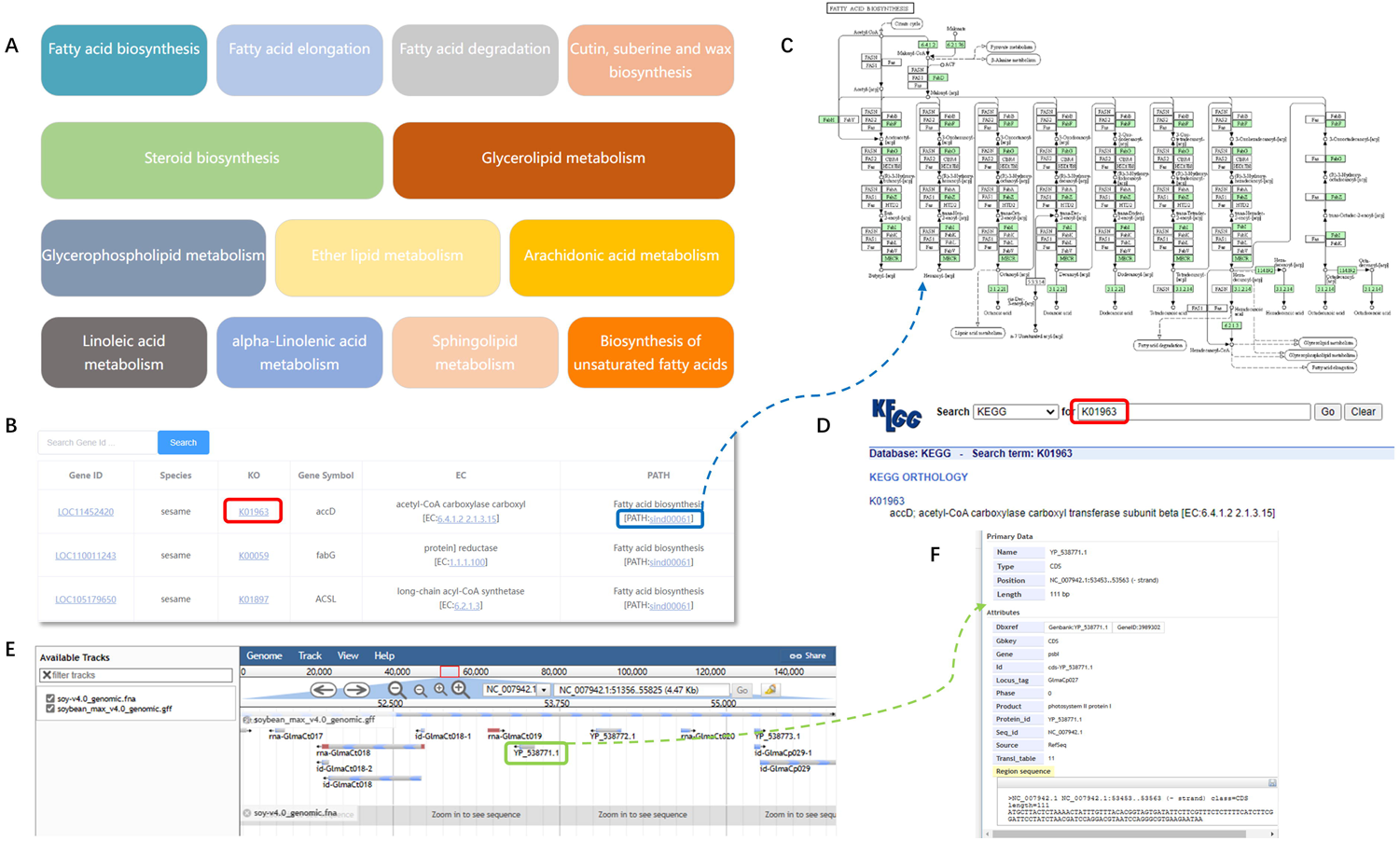
Result of pathway enrichment for lipid metabolism genes in oilseed crops. (A) Lipid metabolism genes from all oilseed crops were enriched for relevant pathways based on the classification criteria of the KEGG database. (B) Information on the gene list for each species displayed after clicking on each pathway. (C) Diagram of the KEGG pathway corresponding to the enrichment result. (D) KEGG term information corresponding to the annotation result of a specific gene. (E) The visualization result for a specific gene in the genome browser. (F) Detailed information displayed after clicking on a selected region in the genome browser.

#### 2. Construction of expression profiles and co-expression networks

Based on the integration of lipid metabolism-related gene sets, we used CLAIR to conduct further analyses to provide more comprehensive information about the temporal characteristics and regulatory mechanisms of gene expression during seed development. This involved constructing complete expression profiles and co-expression networks for each oilseed species, utilizing data from nearly 2,000 batches of lipid metabolism-related transcriptomes (Figure 3A). The “Expression” module on the home page of each oilseed species presents the results of this analysis. The global expression profiles of each species are displayed as heatmaps (Figure 3B). The horizontal coordinates represent the sample numbers of each BioProject in the public database, and the vertical coordinates contain relevant links to detailed information about each BioProject and detailed pathway information (Figure 3C).

**Figure 3.**
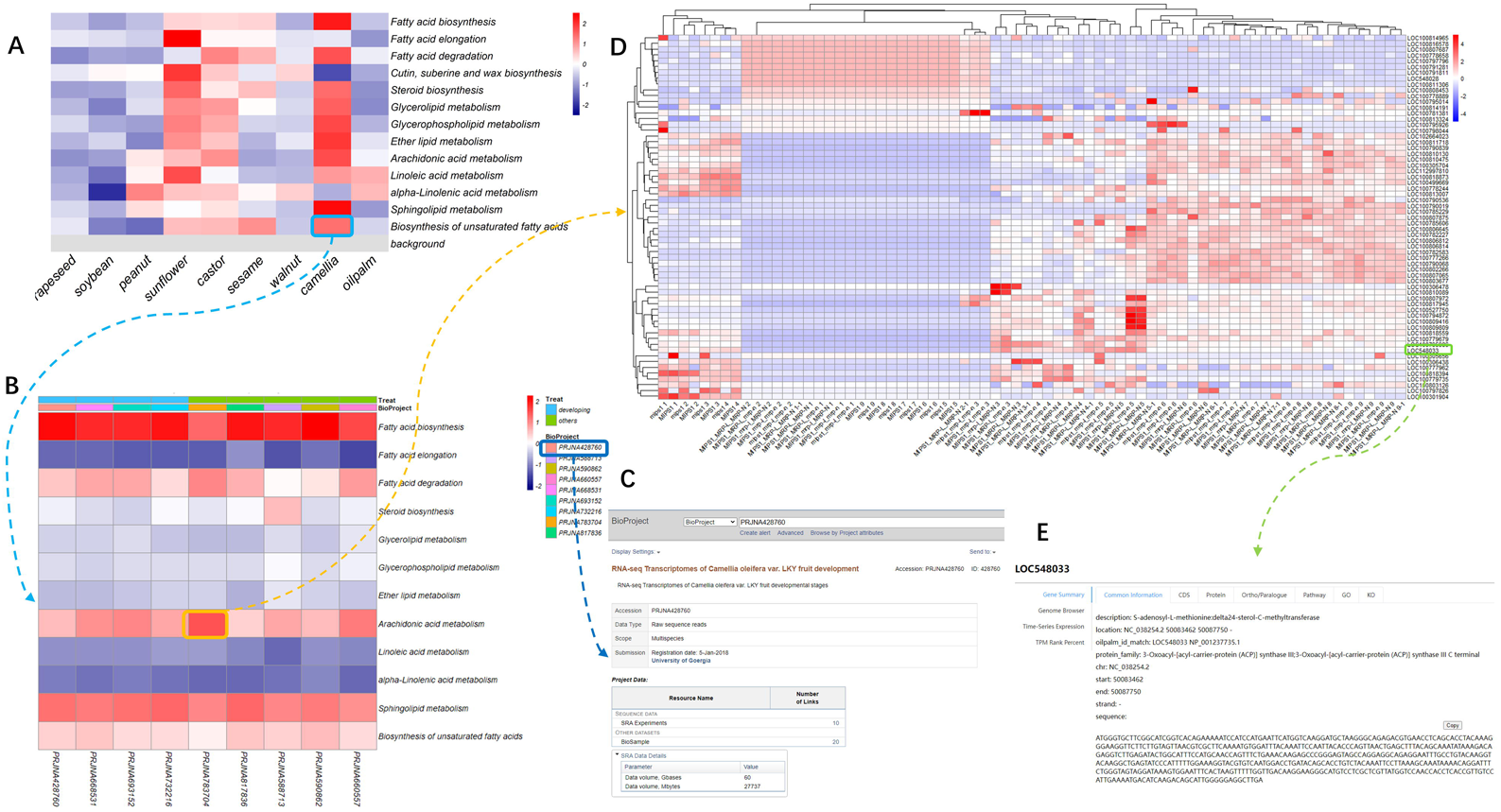
Expression atlas of lipid metabolism related genes in oilseed species based on bulk data. (A) Expression of different lipid metabolism related pathways in nine oilseed species, where the vertical coordinate is the pathway type based on the classification results of the KEGG database, where the background refers to the result of homogenization of the total expression of genes other than lipid metabolism genes. (B) Expression profiles of an oilseed species involved in metabolism related to seed oil content, with all relevant BioProject numbers in the horizontal coordinates. (C) Detailed information about each data set is available by clicking on the BioProject number. (D) Each colored block in Figure 3B generates a new heatmap when clicked, where the vertical coordinates are all genes of the pathway and the horizontal coordinates are the labeling information of all samples under the BioProject, which we have manually organized. (E) Clicking on the gene ID in the vertical coordinate of Figure 3D will again jump to the “common information” interface corresponding to that gene.

To ensure reliable comparisons between transcriptomes, we transformed gene expression levels into SPM format to mitigate the batch effect. Each color block within the global expression profile is clickable and generates an independent heatmap when clicked. The horizontal coordinates of the independent heatmap display manually collated information from all samples within this BioProject, and the vertical coordinates show the IDs of all genes associated with the corresponding pathway of that color block (Figure 3D). Clicking on a specific gene ID allows users to access the gene set section for more information about the gene of interest (Figure 3E). Additionally, users have the option to perform searches based on a target gene ID. The search results will display the module in which the gene of interest is located, and clicking on the module will reveal all the genes within that module. This functionality enables users to comprehensively explore the relationships between genes, pathways, and expression profiles.

Co-expression networks are often used to elucidate functional relationships and regulatory mechanisms between genes. These networks define highly connected gene clusters as modules that are collectively involved in specific biological processes, with kME values often serving as a measure of the relationship between genes and modules (Serin et al., 2016). Based on the results of pre-tagged data classification, CLAIR selected the expression matrix of the data batch with the smallest batch effect in each oilseed species for co-expression network construction and extraction of lipid metabolism-related gene modules. Most of the lipid metabolism genes were evenly distributed across several modules. By clicking on a module, all genes within that module are displayed, and clicking on each gene ID again leads to the “Common information” module, which displays all relevant information for that specific gene. Furthermore, to account for the differential expression of lipid metabolism genes in oilseed crops during development, all transcriptome data involved in seed development are filtered and visually presented for researchers’ convenience.

#### 3. Tools and Resources Guide

CLAIR currently offers three built-in companion tools for the user: JBrowse, BLAST, and ID converter. JBrowse enables users to browse and query information such as oil metabolism genes and transcripts related to oilseed crops, with the Arabidopsis genome displayed by default. This allows users to observe and compare the structures of homologous genes from different oilseed crops. BLAST provides the reference genomes, protein sequences, and nucleotide sequences of all lipid metabolism-related genes from Arabidopsis and the nine oilseed species for comparison. Users can analyze and download sequence files by uploading their own data. However, it is important to note that the lipid metabolism gene sets were constructed based on a specific version of the reference genome, which could be inconvenient to users relying on different versions of genomes. To address this issue, the ID converter integrates lipid-related gene IDs from the genomes of each oilseed species and ID information from popular databases. Users can enter the names of genes, transcripts, proteins, information from SwissProt, TrEMBL, KO, EC, pathways, GO, and PF accessions in various formats based on their needs. Moreover, the ID converter supports gene IDs and protein IDs from different versions of genomes within the same oilseed species. This allows users to easily access and convert relevant gene information, regardless of the reference genome version used.

In addition, the QTL module in CLAIR compiles existing QTL results related to oil content in major oilseed species. By clicking on the corresponding SOC (seed oil content) number, users can access detailed information, including the title of the study, version of the reference genome analyzed, research organization, location data of the identified QTLs, associated gene IDs, and other relevant details. Specifically, this module currently includes information from 11 studies with 215 QTLs in rapeseed, 15 studies with 513 QTLs in soybean, and two studies with 66 QTLs in peanut. These web pages will be continuously updated as new data become available.

## Discussion and future plans

Increasing the oil contents of oilseed crops to meet the growing demand for vegetable oil is a critical objective in modern agriculture (Naylor, 2016; Ortiz et al., 2020). Although several important studies have been conducted, there is currently no comprehensive, up-to-date online database that integrates this information and provides researchers with a wealth of resources related to plant lipid metabolism. In this study, we addressed this gap by constructing the most comprehensive lipid metabolism-related gene set to date for the model plant Arabidopsis. This gene set served as a critical reference to build CLAIR, a user-friendly, resource-integrated database for exploring lipid metabolism-related genes in major oilseed crops based on the extensive literature. CLAIR’s user interface is designed to be concise and convenient for researchers, facilitating data analysis and usage. CLAIR can be used to query lipid metabolism related genes for each species, providing information on protein families, pathways, sequences, expression trends, gene co-expression, and associated QTLs. Moreover, users can download all related data files and utilize practical tools such as GBrowse, BLAST, and ID converter. We plan to expand the content of CLAIR by incorporating information and structures related to plant lipid-related metabolites from MetaCyc, PlantFA, and other databases, thereby increasing the utility of the database and the breadth of information available to researchers.

Oilseed species vary in their oil production periods, geographic distribution, and major oil types (Baud and Lepiniec, 2010). Understanding the underlying causes of these differences serves as a foundation for research aimed at improving oil content in crops. Evolutionary processes, such as contractions and expansions of genes involved in lipid metabolism, may have played a role in shaping the diversity of oil types and oil content among different species. Changes in gene families also affect genome size, thereby influencing metabolic pathways and regulatory networks, which in turn may have implications for organismal adaptation and evolution (Canestro et al., 2007). Using CLAIR, researchers can now gain clearer insights into the overall expression trends of genes within relevant modules by acquiring more complete gene sets, expression profiles, and co-expression networks from existing oilseed species. Besides, by analyzing the differential expression characteristics of genes at high resolution, it will be possible to identify genes of interest for further investigation (Gerstein et al., 2014). Subsequent studies will focus on species-specific genes, pathways, and regulatory mechanisms to better understand the factors contributing to variation in lipid metabolism among oilseed crops.

## DATA AVAILABILITY

CLAIR is available online for free at http://www.clipair.cn/ and does not require user registration.

